# Do you have enough space? Habitat selection of insectivorous cave-dwelling bats in fragmented landscapes of Eastern Amazon

**DOI:** 10.1101/2023.12.07.570662

**Authors:** Valéria da C. Tavares, Mariane S. Ribeiro, Xavier Prous, Alice Araújo Notini, Nathalia Y. Kaku-Oliveira, Leandro M. D. Maciel, Sérgio Sales, Juliana M. Longo, Flávia M. Evangelista, Lucas Rabelo, Iuri V. Brandi, Santelmo Vasconcelos, Sonia Talamoni, Guilherme Oliveira, Leonardo C. Trevelin

## Abstract

Individual movements of bats result from a compromise between their recognition of the environment and their potential to fulfill bats’ life requirements, and to the potential threats associated with moving, all of this is mediated by habitat selection. Mining activities produce modifications to the environments that add heterogeneity and fragmentation to the landscapes used by bats, with overall poorly understood consequences to their movements and to the underground-related biodiversity. Cave dwelling bats spend a large part of their life cycle within their roosts, and they are of paramount importance to the subterranean biodiversity because of their constant movements between the external landscapes, which they selectively use, and the underground ecosystems, where they usually apport energy in form of organic matter. We investigated the variation of patterns of habitat use and selection by cave-dwelling bats in a mosaic of disturbed and conserved Eastern Amazonian forests and rupestrian iron-rich savannas (cangas) interspersed in an extensive iron-caves system. We studied the movements of two phylogenetically distant related insectivores, the aerial insect-catcher *Furipterus horrens* (Furipteridae) and the foliage gleaning bat *Lonchorhina aurita* (Phyllostomidae), both cave-dependent in the study area, one of them categorized as threatened to extinction in Brazil, and virtually unknown in terms of their movement behavior, and habitat use. We used radio telemetry to assess habitat use, under the prediction that these species prefer natural and conserved habitats for their foraging bouts, avoiding human-modified habitats. We also anticipated larger range-areas and commuting distances for both species when foraging in disturbed landscapes. Thirty-one bats were monitored in different landscapes (conserved Amazonian ombrophilous forests and cangas, mining sites and pasture) resulting in an average range of 346.9 ± 762.5 ha and an average commuting distance of 1921.5 ± 2269.7 m for *F. horrens* and of 716.8 ± 1000.6 ha and 2582.2 ± 1966.0 m for *L. aurita*. Our habitat selection analyses suggest that *Furipterus horrens* is an open space forager, with individuals frequently recorded foraging in cangas, and *L. aurita* is a forest forager, although using all habitats proportionally to their availability in the landscape. We did not detect landscape-related effects to the use of space by both species, whereas seasonal variation emerged as a relevant effect. This is the first time that movement data for *F. horrens* and *L. aurita* is presented. Our study delivers original baseline data on movement behavior and conservation of these threatened cave-dwelling bat species with virtually unknown biology and shed light into constraints related to the optimal and adjusted biological cycles of the bats and their range areas under scenarios of disturbance.

## Introduction

The behavior displayed by an individual in the wild is a compromise between sets of possible responses enabling it to cope with adverse situations, such as those arising from anthropogenic activities [1]. Movement behaviors represent conciliations between exploring the environment to meet life requirements and coping with all potential threats associated [2, 3]. Those trade-offs are mediated by habitat selection according to which, regarding to their use of space, the individuals and species optimize their fitness. [4]. Understanding habitat selection may shed light to behavioral mechanisms that allow species to exploit heterogeneous environments [5] and provides insights about the influences of habitat quality on the individuals and populations [1].

Cave-dwelling bats spend roughly half of their life cycles roosting underground, for resting, mating, and raising offspring [6, 7], and the other half searching for, and gathering resources during night foraging bouts in the external surrounding landscapes [8]. Energetic requirements of bats, and the heterogeneity of the landscapes are among the main limiting factors shaping these nightly outside hunts driven by habitat selection and related to the bat’s movements [9]. The interplay between the underground and the surrounding landscapes shapes overall biodiversity patterns in subterranean environments worldwide [7].

Although caves have been usually a focus for the conservation of the cave-related biota [10] the surrounding landscapes can represent most of the home range of an individual bat, playing a major role in the maintenance of the diversity and abundance of bats through time [11]. This is also true in the case of subterranean invertebrate assemblages, which are mostly supported based on bat guano deposition [8, 12].

The conservation of subterranean ecosystems constitutes a modern challenge, given their unique biodiversity that often spatially coincide with the clustered nature of mineral resources [13] often concentrated in tropical areas. Mining activities result in intense, multi-scale landscape modification, and their consequences to the underground biodiversity are poorly understood demanding innovative studies to unveiling biological patterns of these systems and to provide indicators to, ultimately, convey agile and reliable insights to monitoring programs [14, 15, 16]. Understanding and monitoring patterns of habitat selection by cave-dwelling bats potentially provide timely behavioral evidence of changes before worse scenarios of impact take place [1], and that is key to the conservation and management planning of the bats and to the associated cave biodiversity [8].

Despite of the crucial value, for bat conservation, of understanding how bat movements are affected by habitat loss and fragmentation e.g. [17, 18] data on the use of space, habitat preferences and selection by bats have seldomly been assessed in the Neotropics e.g. [19, 20, 21, 22, 23]. Information is even scarcer for Amazonian bats and virtually no data is available for bats using tropical cave systems. This is particularly critical for karstic landscapes demanding management actions for habitat protection, such as the unique iron cave complexes from Eastern South America.

One of the most extensive iron cave systems in the world is nested in the Eastern Amazonia region called “Serra dos Carajás” primarily enclosed within the Carajás National Forest that contains a mosaic of conserved ferruginous canga and Amazon forests, and iron ore exploitation areas, harboring a rich cave bat fauna [24]. Some of these species are regionally cave-dependent, such as the phylogenetically distant and morpho-ecologically highly distinct insect-feeding Thumbless bat *Furipterus horrens* (Chiroptera: Furipteridae) and the Common Sword-nosed bat *Lonchorrina aurita* (Chiroptera: Phylostomidae).

Thumbless bats *Furipterus horrens* Bonaparte, 1837 are tiny bats (3–4 g) with a delicate appearance, broad wings and a long tail extending for most of their large uropatagium, and small thumbs enclosed by the wing membrane. Populations of *Furipterus horrens* are generally associated to humid forested areas and limestone rocky shelters, where they form from small groups to large colonies in Venezuela, in the transition of semideciduous forests and the caatinga, and in the caatinga of Southern and Northeastern Brazil [25, 26] in rock outcrops in the riverbed of the river Xingu [27] and or small associations of one to two bats, one always juvenile in fallen trees of Paracou [28]. Nonetheless, in Carajás, this species relies heavily on the iron caves complexes, where large populations live [24]. The biology of *F. horrens*, a species considered as one of the rarest Neotropical bats, is scarcely known [26]. Most of its South American distribution is included within Brazil, where this species has been listed as “Vulnerable”, according to IUCN criteria based on rough estimates of percentage of mature individuals per population, and because of its association with karstic landscapes under the influence of mining activity, cattle ranching, habitat fragmentation and urban expansion [29].

*Lonchorhina aurita* Tomes 1863 (Phyllostomidae) is a medium-sized foliage-gleaning insectivorous bat, categorized as “Near threatened” in Brazil [29] that possess a singular echolocation system considered an adaptation to aerial hawking [30]. This species has been found roosting in tree hollows [31] but as *F. horrens*, it extensively occupies iron caves in Carajás [24]. For both *F. horrens* and *L. aurita*, virtually nothing is known about their use of space and selection of habitats.

The heterogeneous landscapes of Carajás region from Eastern Amazonia concentrate over 1500 iron caves distributed within well conserved forests and Cangas, and in a mosaic of areas disturbed by iron-mining and peri-rural activities. Our two-fold goals in this study are (1) to describe movement data for *F. horrens* and *L. aurita* generating baseline information on their use of space and habitat selection patterns and (2) to estimate their responses to anthropogenic disturbance at landscape levels. We used radiotelemetry to assess patterns of habitat selection and hypothesized that those two species of insect-feeding cave bats, considered as vulnerable in a large part of their distribution, would rely mainly on well conserved forest and canga habitats to their foraging activities. We predicted the avoidance of anthropogenic, modified habitats by *F. horrens* and *L. aurita*, such as mining and pastures while foraging [10] and that the range areas and commuting distances would be larger at disturbed landscapes for both species.

## Methods

### Study site

We studied the movements of *F. horrens* and *L. aurita* in the National Forest of Carajás (FLONA Carajás), and in the Campos Ferrugíneos National Park, two contiguous Brazilian conservation units located in the Carajás mountain range of southeastern Amazonia, state of Pará, Brazil. Carajás encompasses complex subterranean ecosystems with over 1500 iron caves already prospected [32]. Owe to its vast and unique iron-rich terrains it has also been considered one of the world’s largest deposits of high-grade iron ore [33]. The region is composed of medium altitude plateaus (between 500 and 700 m) scattered along lowlands and forming a mosaic of typical Amazonian rainforests and rocky outcrops with a savannah-like cover, the “canga”. This savannah-like ecosystems are however uniquely adapted to iron-rich soils and dissimilar to other Amazonian savannas, and it is called the “cangas” [32, 34, 35]. The Carajás region currently combines pristine and recovering protected areas and areas for ore exploitation (iron, copper, manganese) and other unprotected areas under anthropogenic use such as pastures, abandoned crops and peri-urban zones [36] – Fig 1). The regional climate is defined as Aw following Köppen classification, marked by high annual rainfall with a well-defined dry season [37].

**Fig 1.**
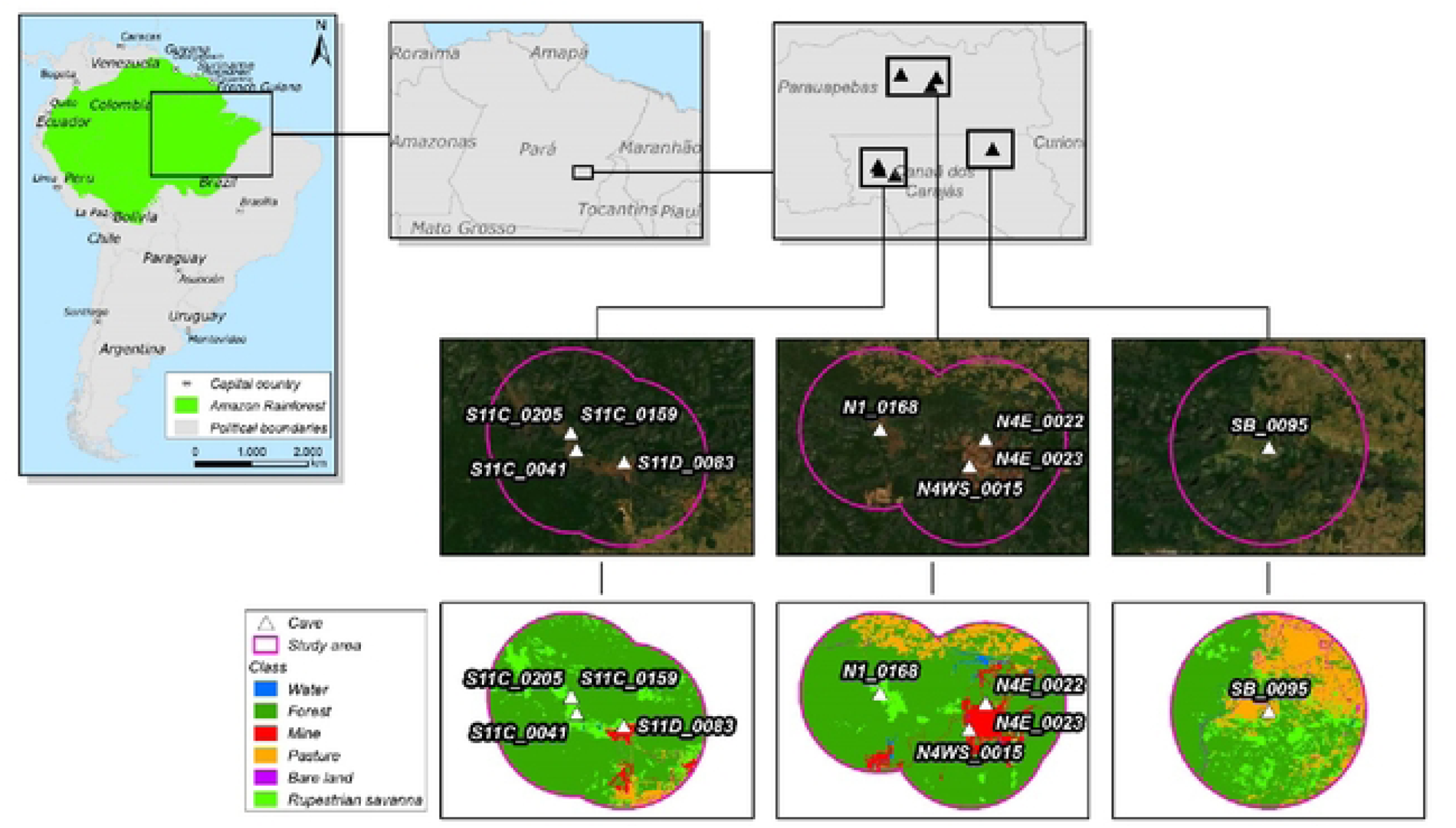
Location of the study area, the Floresta Nacional de Carajás, and study sites containing the caves sampled and the habitats considered in the study, present in the surrounding landscapes.

## Experimental design and data collection

### Landscape data

The composition of the landscapes surrounding each cave sampled was initially accessed within a circular 12km-radius buffer, centered around the main cave entrance, using photo-interpreted data from Sentinel 2 images with 10 meters of spatial resolution. We initially discriminated six habitat types as landscape categories (Forest, canga, mining land, pasture, exposed bare land and water covered land) and quantified their availability within each buffer. Our experimental design was then composed of three categories of landscape dominance (treatments): landscapes dominated by conserved natural habitats (forests, savannah-like “canga”), landscapes dominated by mining land use (mine) and landscapes dominated by pastures (pasture) (Fig 1). All imagery analyses were conducted using QGIS 3.14. We then selected caves for our study based on the combination of two criteria, the representativeness of caves’ surrounding landscapes in the region together with the presence of our target species (*F. horrens* and *L. aurita*).

### Capturing and marking bats

Bats were captured during daytime while roosting inside caves using entomological hoop nets [38], as we wanted to monitor the movements of cave residents. The individuals monitored for each species were treated as replicates of their populations, and each population was sampled twice, one in the wet and one in the dry season. Captured individuals were kept in cotton bags for biometry, age and reproductive assessments [39] and for further tagging with radio transmitters with unique signaling frequencies (LB-2x – Holohil Inc., Canada).

We carefully trimmed patches of upper middorsal fur, close to the scapula horizontal plane (in relation to a longitudinal body axis) of all individuals selected to receive radio transmitters, to attach the radio transmitters with the help of surgical adhesive (Fig 2). The weight of each transmitter was 0.31 to 0.41 g corresponding to roughly 8% of the smaller bat’s weight. Radio-tagged bats were then released inside the caves in which they were captured, and radiotelemetry data begin to be recorded departing from the next night, to avoid bias resulting from the capture and handling of individuals.

**Fig 2.**
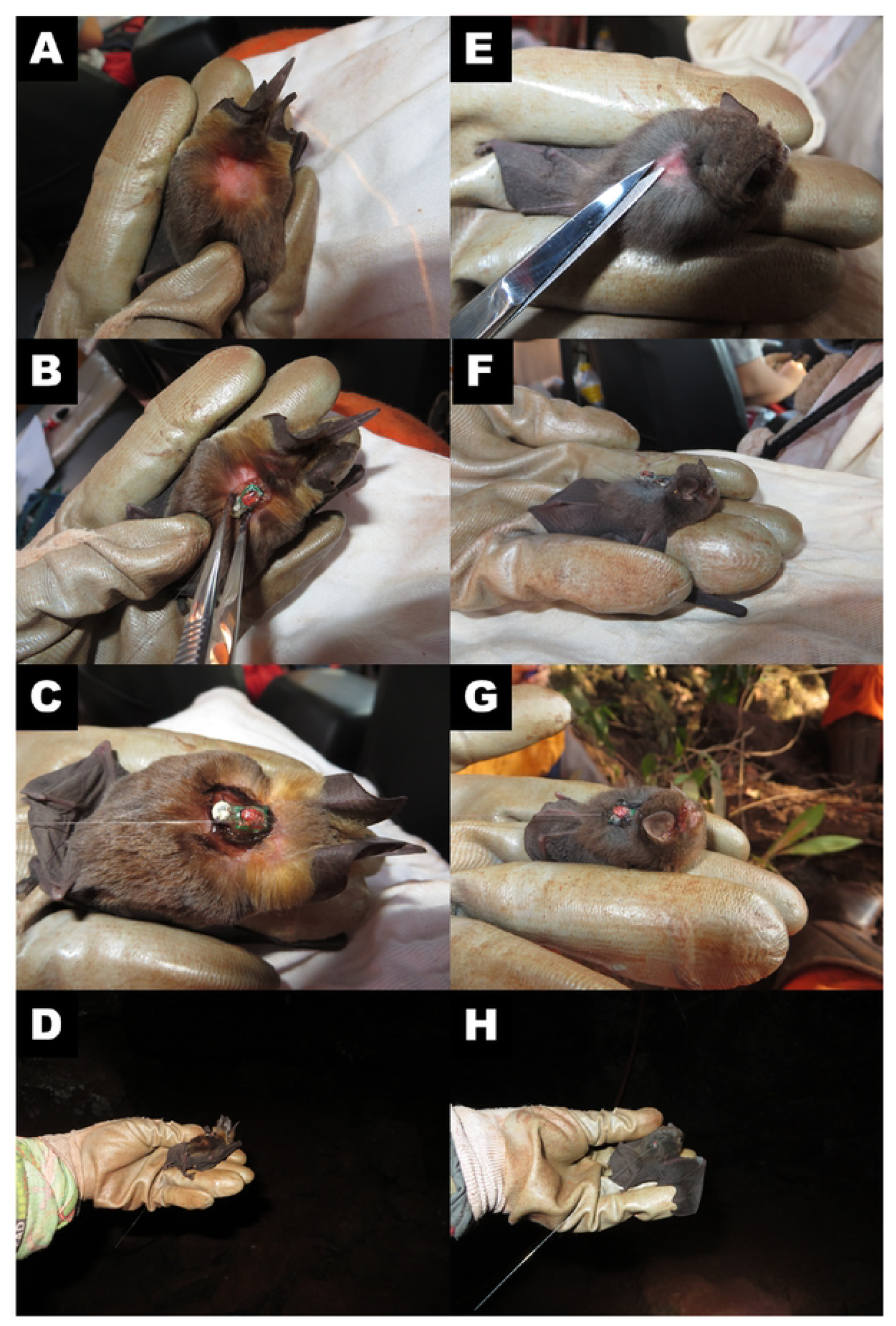
Attachment of radio transmitters, after the clipping of the fur, and release of the bats: A) Hair clipping in the back of a *L. aurita* individual; B) gluing of the transmitter to the back of a *L. aurita* individual; C) Transmitter glued on and D) hand release. E) Hair clipping in the back of a *F. horrens* individual; B) gluing of the transmitter to the back of a *F. horrens* individual; C) Transmitter glued on, and D) hand release.

Only healthy adult and non-reproductive females, as diagnosed by physical examination was selected to receive radio transmitters. We complied with the guidelines from the American Society of Mammalogists [40] for the use of wildlife in research, and this study was authorized by the Instituto Brasileiro do Meio Ambiente e dos Recursos Naturais Renováveis (IBAMA) (ABIO N° 1000/2018).

### Radiotracking

We conducted six sampling sessions, three in the dry season (August and September 2018, July 2019) and three in the wet season (November 2018, February and November 2019). We marked and monitored a total of 46 individuals, including 21 *F. horrens* and 25 *L. aurita* with a similar number of sampled individuals between seasons (see Table 1). A five-day sampling protocol was followed for each marked individual; each sampling night was sub-divided into three 4-h intervals, starting at 18 h and ending at 06 h of the next day. We aimed to sample all intervals for each individual along the five-days sampling periods allowing inferences regarding habitat use in situations where the number of location fixes differs among individuals (replicates) [20, 41]. Signal monitoring was conducted by independent teams, performing active radio signal searches and angulation techniques to obtain locations [42] using radio-receiver model TRX-1000 (Wildlife Materials Inc.) coupled to three-elements Yagi antennas.

Successive locations and roost fixes were mapped to obtain flight routes and range areas.

**Table 1.** Summary of results obtained per species. Species/Individuals, Cave, Treatment, Date, Season, Days (sampled) and N (Number of locations estimated). MC (ha) – Minimal Convex Polygon estimate. KF 95% (ha) – Fixed Kernel estimates of range use with 95% (ha) of samples; KF 50% (ha) – Fixed Kernel estimates of range use with 50% (ha) of samples; MC 95% (m) – Maximum linear distance travelled from cave entrance in KF 95%; MC 50% (m) – Maximum linear distance travelled from the cave entrance to core foraging areas in KF 50%. At the bottom of each species, we present the average and standard deviation of estimates for each “use of space” parameter.

| Species/ Individuals | Cave | Treatment | Date | Season | Days | N | MCP (ha) | KF 95% (ha) | KF 50% (ha) | MC 95% | MC 50% |
| --- | --- | --- | --- | --- | --- | --- | --- | --- | --- | --- | --- |
| *Furipterus horrens* |  |  |  |  |  |  |  |  |  |  |  |
| 1 | SB_0095 | Pasture | out/18 | Dry | 1 | 28 | 27,1 | 55,0 | 8,8 | 1037,6 | 227,7 |
| 2 | N4WS_0015 | Mining | nov/18 | Wet | 5 | 107 | 30,9 | 31,5 | 6,3 | 710,5 | 362,4 |
| 3 | SB_0095 | Pasture | nov/18 | Wet | 5 | 43 | 37,8 | 60,8 | 8,7 | 2002,4 | 275,2 |
| 4 | SB_0095 | Pasture | nov/18 | Wet | 3 | 63 | 39,5 | 74,1 | 9,7 | 1281,8 | 327 |
| 5 | SB_0095 | Pasture | set/18 | Dry | 3 | 35 | 4,3 | 404,2 | 64,8 | 1691,6 | 510,6 |
| 6 | SB_0095 | Pasture | nov/18 | Wet | 5 | 4 | 0,4 | 12,1 | 2,9 | 365,5 | 254 |
| 7 | S11C_0205 | Preserved | dez/19 | Wet | 2 | 8 | 35,2 | 157,4 | 44,1 | 1059,6 | 833,5 |
| 8 | N4WS_0015 | Mining | nov/18 | Wet | 2 | 57 | 1921,9 | 3545,2 | 500,9 | 10852,3 | 2094,5 |
| 9 | SB_0095 | Pasture | set/18 | Dry | 3 | 23 | 1,1 | 3,0 | 0,5 | 366 | 132,6 |
| 10 | SB_0095 | Pasture | set/18 | Dry | 2 | 22 | 53,8 | 192,4 | 53,7 | 1239,5 | 846,2 |
| 11 | S11C_0041 | Preserved | out/18 | Dry | 2 | 6 | 35,7 | 387,9 | 76,0 | 2558,3 | 1063 |
| 12 | S11C_0041 | Preserved | out/18 | Dry | 5 | 60 | 38,0 | 72,3 | 18,3 | 1201,6 | 878,5 |
| 13 | N4WS_0015 | Mining | dez/18 | Wet | 4 | 29 | 126,8 | 399,4 | 97,2 | 2515 | 1183,8 |
| 14 | SB_0095 | Pasture | nov/18 | Wet | 5 | 56 | 263,4 | 728,7 | 113,1 | 4352,2 | 763,5 |
| 15 | S11C_0205 | Preserved | dez/19 | Wet | 5 | 112 | 54,3 | 44,5 | 7,0 | 774,4 | 410,8 |
| 16 | N4WS_0015 | Mining | ago/18 | Dry | 2 | 34 | 14,9 | 40,0 | 8,4 | 983 | 845,5 |
| 17 | N4WS_0015 | Mining | ago/18 | Dry | 2 | 30 | 25,6 | 86,4 | 17,3 | 1068 | 394 |
| 18 | N4WS_0015 | Mining | ago/18 | Dry | 2 | 26 | 15,8 | 47,9 | 12,5 | 936 | 763,8 |
| 19 | N4WS_0015 | Mining | ago/18 | Dry | 3 | 21 | 25,7 | 96,0 | 25,5 | 965 | 772 |
| 20 | S11C_0041 | Preserved | set/18 | Dry | 9 | 62 | 309,0 | 670,4 | 155,8 | 3283,9 | 2630,3 |
| 21 | S11C_0205 | Preserved | dez/19 | Wet | 3 | 11 | 35,9 | 175,7 | 52,0 | 1107 | 812 |
| Mean ± SD |  |  |  |  |  |  | 147.5 ± 414.5 | 346.9 ± 762.5 | 61.13 ± 109.2 | 1921.5 ± 2269.7 | 780.0 ± 608.7 |
| *Lonchorhina aurita* |  |  |  |  |  |  |  |  |  |  |  |
| 1 | N4WS_0015 | Mining | ago/18 | Dry | 4 | 10 | 19,7 | 93,5 | 22,1 | 1345,4 | 1169,2 |
| 2 | N4WS_0015 | Mining | ago/18 | Dry | 6 | 48 | 111,1 | 163,1 | 21,5 | 1982 | 483,5 |
| 3 | N4WS_0015 | Mining | ago/18 | Dry | 4 | 5 | 32,7 | 359,1 | 88,0 | 2079 | 917,6 |
| 4 | N4WS_0015 | Mining | ago/18 | Dry | 4 | 19 | 80,2 | 196,9 | 53,6 | 1568 | 1280 |
| 5 | S11C_0041 | Preserved | dez/19 | Wet | 2 | 3 | 3,9 | 103,8 | 27,6 | 846,6 | 611,2 |
| 6 | N4WS_0015 | Mining | fev/19 | Wet | 5 | 12 | 160,2 | 566,3 | 125,6 | 2168,2 | 1543,7 |
| 7 | S11C_0041 | Preserved | nov/19 | Wet | 4 | 6 | 413,1 | 2495,8 | 620,5 | 4185,9 | 2690,7 |
| 8 | N4WS_0015 | Mining | ago/18 | Dry | 4 | 43 | 123,0 | 210,0 | 42,8 | 1883,6 | 1530,5 |
| 9 | S11C_0041 | Preserved | nov/19 | Wet | 5 | 7 | 113,3 | 602,5 | 145,7 | 2130,2 | 890,1 |
| 10 | S11C_0041 | Preserved | jul/19 | Dry | 1 | 7 | 27,3 | 197,3 | 55,3 | 1659 | 1314,3 |
| 11 | N4WS_0015 | Mining | fev/19 | Wet | 3 | 20 | 47,3 | 155,0 | 42,7 | 1371 | 961,4 |
| 12 | N4WS_0015 | Mining | ago/18 | Dry | 4 | 14 | 40,3 | 177,6 | 50,4 | 1651,3 | 1384,6 |
| 13 | SB_0095 | Pasture | nov/19 | Wet | 2 | 4 | 8,3 | 291,3 | 74,6 | 1658,7 | 1100 |
| 14 | SB_0095 | Pasture | set/18 | Dry | 2 | 3 | 22,6 | 311,2 | 87,6 | 1945,3 | 1536 |
| 15 | S11C_0041 | Preserved | jul/19 | Dry | 3 | 3 | 29,2 | 715,0 | 189,2 | 1938,9 | 1355,2 |
| 16 | SB_0095 | Pasture | set/18 | Dry | 4 | 6 | 226,0 | 2326,3 | 499,0 | 5149,5 | 1614,3 |
| 17 | SB_0095 | Pasture | nov/19 | Wet | 1 | 5 | 7,1 | 631,7 | 169,7 | 1692,2 | 1192 |
| 18 | SB_0095 | Pasture | set/18 | Dry | 2 | 35 | 54,7 | 105,1 | 21,7 | 1463,5 | 872,3 |
| 19 | SB_0095 | Pasture | nov/19 | Wet | 4 | 32 | 4291,8 | 3947,5 | 514,5 | 9950,3 | 4663,7 |
| 20 | SB_0095 | Pasture | set/18 | Dry | 6 | 44 | 1259,5 | 2579,0 | 483,9 | 5148,6 | 3826,5 |
| 21 | S11C_0041 | Preserved | nov/19 | Wet | 3 | 27 | 28,2 | 65,1 | 12,2 | 1268,8 | 422,5 |
| 22 | SB_0095 | Pasture | set/18 | Dry | 3 | 23 | 39,8 | 122,2 | 28,2 | 2352 | 1713,9 |
| 23 | S11C_0041 | Preserved | nov/19 | Wet | 5 | 9 | 101,8 | 412,4 | 109,2 | 1718 | 1020,9 |
| 24 | N4WS_0015 | Mining | fev/19 | Wet | 5 | 41 | 450,9 | 741,1 | 133,2 | 5069,7 | 2293,4 |
| 25 | S11C_0041 | Preserved | jul/19 | Dry | 1 | 9 | 84,8 | 351,4 | 98,7 | 2329,8 | 1579,8 |
| Mean ± SD |  |  |  |  |  |  | 311.1 ± 868.7 | 716.8 ± 1000.6 | 148.7 ± 177.6 | 2582.2 ± 1966.0 | 1518.7 ± 970.8 |

## Data Analysis

### Home range estimates

We used Best Bi-angulation Estimators to determine locations, after correcting for magnetic declination in all compass bearings, using LOAS™ (Ecological Software Solutions, Inc.). Location datasets determined for each individual were incorporated into QGIS 3.14 software and subsequently, range areas were estimated using the HRT 2 extension, Home Range Tools [43].

We generated range area estimates for each individual following Kernohan *et al*. [44], as the ‘‘extent of area with a defined probability of occurrence of an animal during a specified time period’’. We used fixed normal kernel density estimations (KF) with 95% (KF 95%) and 50% (KF 50%) kernel isopleths to delineate the range areas [45], using least-squares cross-validation (LSCV) to estimate the smoothing parameter (H) [46]. In addition to KF 95% and KF 50%, we generated range area estimates using Minimum Convex Polygon (MCPs) estimates, uniting the most external locations.

These range estimates were subsequently used in our hypotheses testing.

### Habitat use/selection

To evaluate habitat selection patterns, we estimated individual-based habitat uses by contrasting the proportional use of habitats by each individual with our quantified amounts of habitat availability. Since “habitat” is a scale-dependent concept we organize our data following Johnson’s [47] hierarchical orders of habitat selection. In our framework, the second-order selection represents the proportional habitat use considering 95% kernel density estimations (KF 95%), and the third-order selection is the proportional habitat use considering KF 50% kernel density estimations (KF), both within the MPC estimated for each individual.

We used the non-parametric analytics as proposed by Fattorini et al. [48] based on the combination of sign tests. Within that framework, each habitat is individually tested, and an overall statistic value for the simultaneous assessment of habitat selection in all habitat types can be obtained by combining P values from each test through permutation.

We considered all six habitat types (Forest, canga, mining land, pasture, exposed bare land and water covered land) and assessed whether each habitat was proportionally used according to its availability for each bat or whether it was over or under-utilized. All analytical procedures were conducted within the R environment [49], using package “phuassess” for R 3.3.1 [50].

### Modelling use of space

To evaluate the effects of landscape variation in the movement behavior of the bats we modelled their use of space using four parameters: range area estimates (KF 95% area in hectares), core-foraging area estimates (KF 50% area in hectares) and two commuting distances, the Maximum linear distance (m), travelled from cave entrance in KF 95%, MC 95%, and Maximum linear distance (m), travelled from the cave entrance to core foraging areas in KF 50%, MC 50%. We implemented linear mixed effect models (LMMs) in a maximum likelihood framework [50] using these parameters as response variables and landscape treatments (conservation/disturbance), seasonality, and their interaction as fixed effects.

We also modelled cave and species as random effects, and data for both species in all the caves were polled for these procedures. We used multi-model inferences to evaluate competing candidate models representing all possible combinations of these variables. Model selection was based on the Akaike Information Criteria (AIC), considering all top-ranking models with ΔAIC < 2 as having similar support [52]. We adopted a stepwise modelling, first selecting between random effects using Restricted Maximum likelihood, then selecting between fixed effects using standard Maximum likelihood, and lastly assessing estimates of the final models [50, 53].

All analyzes were conducted within the R environment [49], using “lmertest” and “MuMIn” packages [54, 55] and all models achieved convergence.

## Results

We successfully monitored seventeen individuals of *F. horrens* and 14 of *L. aurita* in all night intervals and sampled for more than one day or 9 locations. Sampling lasted three and a half days on average for each individual of both species (*F. horrens*: 3.4 ± 1.31; *L. aurita*: 3.6 ± 1.4). We sampled a total of 1168 locations, and the mean number of locations obtained per individual was 40.5 ± 30.3 for *F. horrens* and 17.8 ± 15.1 for *L. aurita*. Initial exploratory analysis did not indicate a significant relationship between the size of estimated range areas and the number of locations obtained for either *F*. *horrens* (r = 0.09; p = 0.74) or *L. aurita* (r = 0.29; p = 0.34). There was also no significant relationship between the estimated range areas and sex for either *F. horrens* (t = 0.79; d.f. 7.88; p = 0.45) or *L. aurita* (t = -0.28; d.f. 3.44; p = 0.79). Our exploratory results corroborated the pooling of individuals as replicates for subsequent analyses, and the results obtained for each individual sampled are summarized in Table 1.

### Habitat use/selection

Our habitat selection estimates for *Furipterus horrens* indicate a trend to foraging in open spaces. By contrasting the KF 95% range area with habitat availability (second-order selection), we observed that individuals of *F. horrens* preferentially used non-forested habitats, mainly the canga and pastures, compared to the proportional use of any other type of habitat, and there is also a pattern of avoidance of forested and water covered habitats (p=0.0004, Fig 3a). The trend to the use of open areas appears only subtly when contrasting the foraging core area range KF50% and MPC habitat availability (third-order selection) but it was not corroborated (p>0.05, Fig 3b).

**Fig 3.**
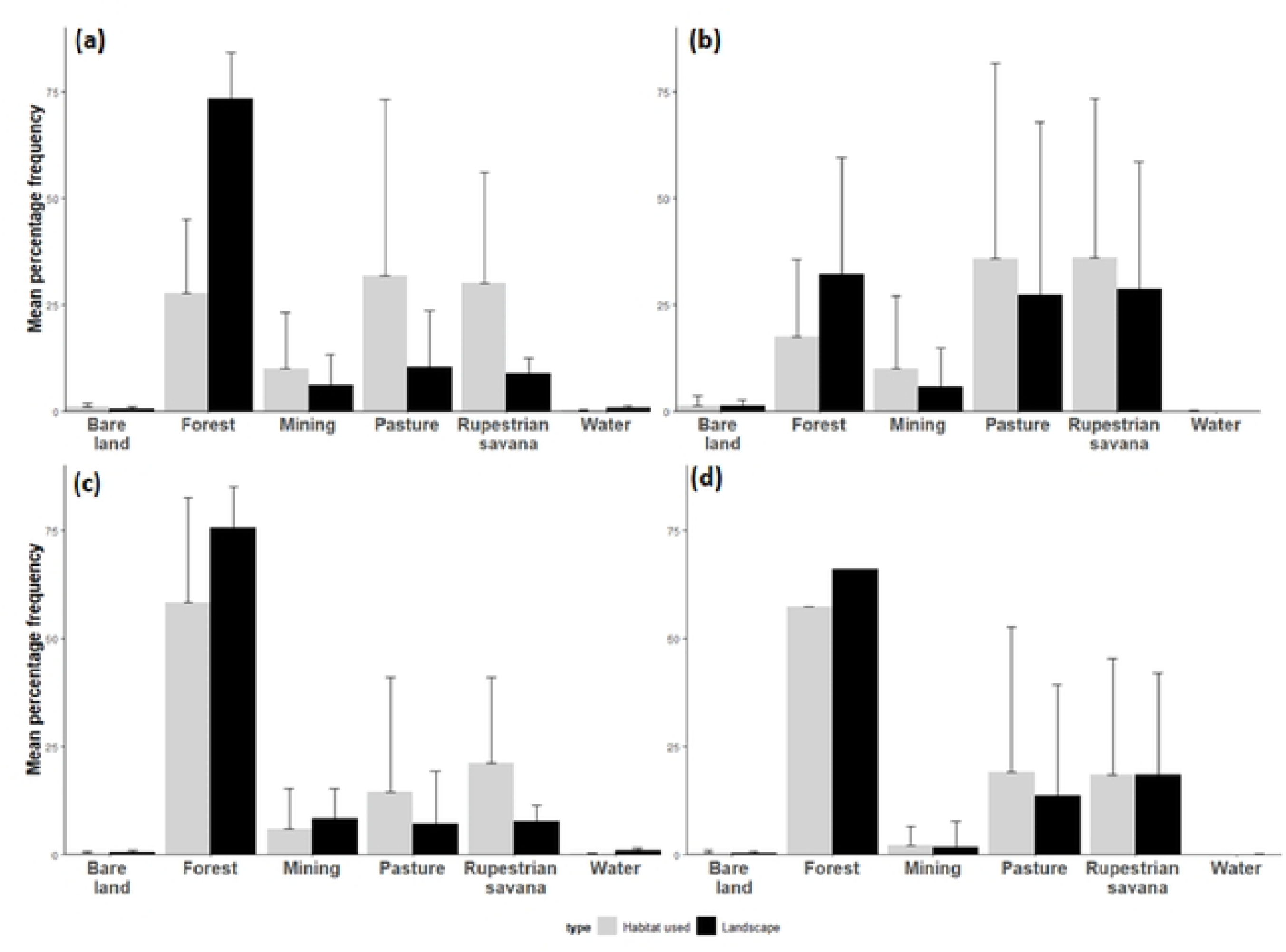
Landscape variation (bare land, forest, mining land, pasture, rupestrian savannah, and water) and habitat availability considering the estimative of KF 95% foraging area. (a) and KF 50% foraging area (b) for *Furipterus horrens*, and KF 95% foraging area and KF 50% foraging area by *Lonchorhina aurita* (c) and (d).

*Lonchorhina aurita*, on the other hand, used all habitats proportionally to their availability in the landscape, except for the avoidance of this species of water covered habitats (Second order selection, p=0.005, Fig 3c). Furthermore, contrasting core foraging areas (KF50% defined) and MPC habitat availability, again *L. aurita* used all habitats proportionally to their availability in the landscapes (Third-order selection, p>0.05, Fig 3d). Forested habitats were the most used by this species and the rupestrian savannahs (cangas) and pastures appear only as complementary (Fig 3d).

### Use of the space

For all adjusted models, cave species were selected as random factors in fixed slope model schemes, and landscape types and seasonality were selected as fixed effects in the final models. Data on the use of disturbed landscapes (cangas and pastures) indicate a trend to accommodate larger foraging areas and longer commuting distances than conserved landscapes use data in the wet season, and the inverse pattern occurs in the dry season (Fig 3). In the range area estimate models, when KF 95% is considered, there was a marginal effect of seasonality (F=3.89; df=2; p=0.059). In contrast, there was no fixed effect narrowing to the core foraging areas (KF 50%) (Fig 4). Likewise, in the commuting distances models considering a maximum KF 95%, the commuting distance was affected by seasonality (F=8.17; df=2; p=0.008) but considering a maximum KF 50%, no fixed effect was found. Overall, our data indicate the influence of seasonality in the landscape use that was not observed in the bat’s core foraging area.

**Fig 4.**
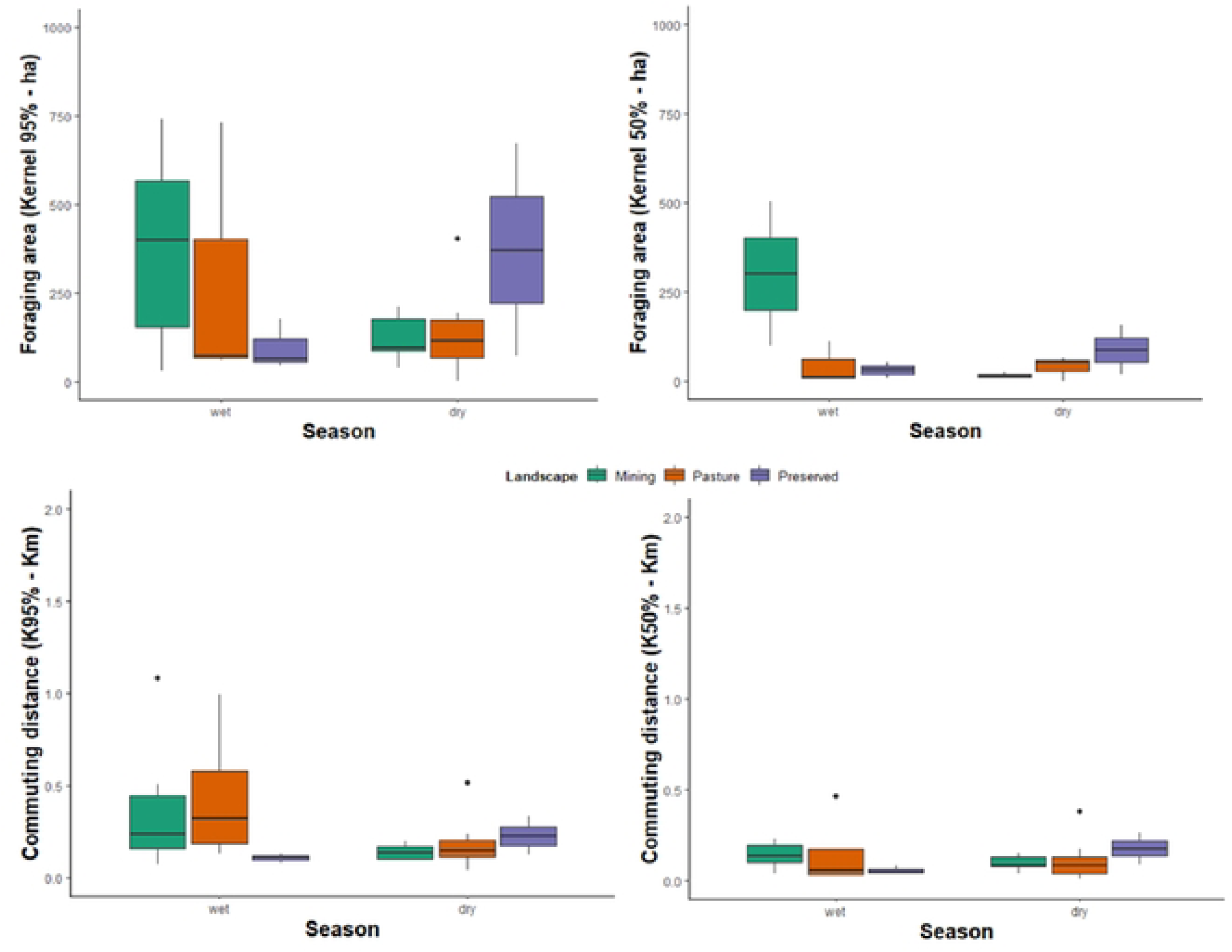
Effects of landscape variation and seasonality for the combined estimative of the foraging areas of *Furipterus horrens* and *Lonchorhina aurita* considering the fixed kernel estimates, KF 95% (a) and the KF 50% (b), and the Maximum commuting distances 95% (c) and 50% (d).

## Discussion

The question to whether the insectivorous, cave-dwelling species *Furipteus horrens* and *Lonchorhina aurita* have enough space to forage in Carajás mosaics has been partially responded by our data. We overall failed to support our predictions that landscape disruption limits the use of space by the two bat species but had indicatives that support our expectations of differential use of habitats by the two bat species studied. We assessed the selection of open areas for *F. horrens* and those included natural, conserved canga habitats, and the open degraded habitats represented by pastures. For the individuals of *L. aurita* that we marked with transmitters and tracked, forests were by far the most used habitats, and the canga and the pastures were complementary foraging areas. A key point to our analysis of habitat preferences was the assumption that certain habitats may be avoided or preferred relatively to others [50] and relative to what is available in the landscape. The sword-nosed bat *L. aurita* used, proportionally, most of what was available of forests, whereas the abundant forested habitats available for *F. horrens* were somewhat neglected by this species. Noteworthy results found for both species were that their use of space is markedly related to seasonality rather than to landscape variation alone.

Data on movement behavior of Neotropical bats in South America is scarce and include mostly estimates for phylostomid species, such as nectar feeding bats from the Brazilian cerrado, *Glossophaga* and *Lonchophylla* [21], fruit bats from the Brazilian Atlantic Forest *Carollia*, *Artibeus*, and *Sturnira* [22], fruit eating, and insect-eating, foliage gleaning bats from Tapajós region, Eastern Amazon *Artibeus*, *Carollia*, *Gardnerycteris, Tonatia,* and *Trachops*, and fish eating bats (Noctilionidae) [23]. To our knowledge, ours are the first estimates of movement behavior for both *L. aurita* (Phyllostomidae), and for the family Furipteridae, as represented by *F. horrens* and that is why we cannot treat them in a broad comparative fashion.

Variations in habitat quality and availability fine-tuned with specific foraging strategies are expected to affect range area estimates, as discussed elsewhere [19, 20, 56, 57]. Overall, our estimates of area range and commuting distances for both species studied are larger, on average, than previous estimates found for other species of Neotropical bats [19, 20, 21, 23]. This is contrary to our expectations particularly in the case of *F. horrens*, which is one of the tiniest Neotropical bats (3 to 4 g) as range sizes are expected to increase with body size and be limited by the energetic constraints in mammals [58, 59] including bats [60].

The biology of *Furipterus horrens* is scarcely known [26, 62], and the little dietary data available recovered some evidence to the consumption of moths [26, 61]. All Neotropical bats use echolocation calls to some extent to navigate and search for food. The echolocation call patterns of *F. horrens* were described in more detail recently revealing that these bats emit short and extremely high frequency and broadband pulses that range from 70 to 210 kHz [62]. Based on the characteristics of the calls of *F. horrens*, resembling the calls of species of Old-World vespertilionid bats from the genus *Kerivoula* [63] Falcão and collaborators [62] suggested that they may also forage in dense vegetation. Our results appear to conflict with this suggestion because the combination of the movement patterns we found with the echolocation behavior described, is suggestive of patterns related to the *Edge Space Aerial foragers* guild as described by Denzinger et al. [64]. Several species of moths have evolved sensitive ears to hearing the bats in one of the clearer examples of evolutionary predator-prey models [65]. Although along this “arms-race” some moths have evolved the capacity of hearing pulses much above 200 kHz, such as the case of greater wax moth, *Galleria mellonella* (Insecta: Lepidoptera) which can hear ultrasonic frequencies near 300 kHz [66] the combination of high frequency, low intensity and short calls of *F. horrens* may enable these bats to overcome the hearing abilities of moths [62], explaining in partially the conflicting results.

*Lonchorhina aurita*, a foliage-gleaning, animalivorous phylostomid has been traditionally considered a narrow space forager and a “forest bat” [67]. As most animalivorous phyllostomids, bats from the genus *Lonchorhina* have been considered passive gleaning foragers relying primarily in signs provided by their prey [64, 68]. However, several studies revealed refinements to our understanding of the evolution of foraging strategies and echolocation patterns of animalivorous phylostomids, including adaptations enabling then to increment their abilities to detect and locate prey with elaborated variations of echolocation calls, and some of them comparable to aerial insectivores from other bat families, such as Emballonuridae and Vespertilionidae (e.g. *Macrophyllum macrophyllum* [55, 69]. A recent study revealed that this is also the case for *L. aurita*, which has an unusual call structure that deviates from the standard call design of phyllostomids bats, perhaps related to its unique noseleaf morphology. The echolocation system of *L. aurita* retains basic phyllostomid characteristics such as multi-harmonic FM-calls, but, as is typical in aerial insectivores, it has evolved to also include narrow-bandwidth CF-components and a terminal group [30]. Specialized foraging patterns among phylostomids have been more and more associated to the modulation of access to target resources and habitats.

Thus, by combining our habitat selection results with the echolocation evidence available we corroborate the suggestion of Gessinger et al. [30] on the evolution of an unusual foraging strategy pattern among phylostomids, in the case of *L. aurita* somewhat like an edge space aerial insectivore notably using primary forested habitats, but also open habitats, accordingly to their availability. Our observations also include patterns of combined use of conserved and disturbed habitats, i.e. movements over fragmented landscapes, and coincide with results obtained in other studies of insectivorous bats [56] and nectar feeding bats from the cerrado [21] and poses further questions related to food constraints for both species, *F. horrens* and *L. aurita*. In central Amazonia, for instance the foraging activity of the cave bat *Pteronotus rubiginosus* was mediated by insect availability, independent of the habitat types (Oliveira et al. 2015). In a few words, it is relevant to investigate how does the current mosaic of conserved and degraded habitats limit the food supply for these species. This is a key question because as movements are costly, food sources are limiting and that is important information for conservation and management.

Food and roosts are key and limiting resources for bats [70] and ultimately, constraints to cave bats will affect cave ecosystems that largely depend upon the guano. This leads to the Brazilian policy concept “area of influence” of the cave, which conveys the idea of the amount of space away from the cave perimeter, measured in kilometers, is needed to safeguard the biodiversity within the cave. Our incipient understanding of the mechanisms affecting the chances of Neotropical cave bats to effectively find and collect food in the surrounding landscapes, and the little knowledge about patterns of roosting selection of Neotropical cave bats prevents us from contributing on estimates on that, and this is crucial for bat and cave ecosystem conservation, and for exercises of prioritization for conservation.

## Acknowledgment

We thank the health and safety teams of companies Ativo Ambiental and Vale S.A. for their help, as well as the technical team of Ativo Ambiental who actively participated in data collection.

## References

1. Romero, LM. Physiological stress in ecology: lessons from biomedical research. Trends in Ecol and Evol. 2004 May;19(5):249–55.

2. Gurarie E, Andrews RD, Laidre KL. A novel method for identifying behavioural changes in animal movement data. Ecology Letters. 2009 May;12:395–408.

3. Mori E, Lovari S, Cozzi F, Gabbrielli C, Giari C, Torniai L et al. Safety or satiety? Spatiotemporal behaviour of a threatened herbivore. Mamm Biol. 2020 February;100:49–61.

4. Rosenzweig ML, Abramsky Z. Centrifugal community organization. Oikos. 1986 May;46:339–348.

5. Schick RS, Loarie SR, Colchero F, Best BD, Boustany A, Conde DA et al., Understanding movement data and movement processes: current and emerging directions. Ecol. Lett. 2008 Dec;11:1338–1350.

6. Trajano E. Protecting caves for the bats or bats for the caves? Chiroptera Neotropical. 1995 Dec;1(2):19–22.

7. Culver DC, Pipan T. The Biology of Caves and Other Subterranean Habitats. Oxford: Oxford University Pres; 2019.

8. 8. Furey NM, Racey PC. Conservation ecology of cave bats. In Voigt C, Kingston T, editors. Bats in the Anthropocene: Conservation of Bats in a Changing World. Springer International Publishing; 2016. p. 463–500.

9. Speakman JR, Thomas DM. Physiological Ecology and energetics of bats. In Kunz TH, Fenton MB, editors. Bat Ecology. Chicago: University of Chicago Press, Illinois; 2003. p. 430–492.

10. Phelps K, Jose R, Labonite M. Correlates of cave-roosting bat diversity as an effective tool to identify priority caves. Biol. Conserv. 2016 Sept 201;201–209.

11. Struebig MJ, Kingston T, Zubaid A, Le Comber SC, Mohd-Adnan A, Turner A et al. Conservation importance of limestone karst outcrops for palaeotropical bats in a fragmented landscape. Biol. Conserv. 2009 Oct 142:2089–2096.

12. Ferreira RL, Martins RP. Trophic Structure and Natural History of Bat Guano Invertebrate Communities with Special Reference to Brazilian Caves. Tropical Zool. 1999 Jul 2:231–252.

13. Jaffé R, Prous X, Zampaulo R, Giannini TC, Imperatriz-Fonseca VL, Maurity C Oliveira G. Reconciling mining with the conservation of cave biodiversity: a quantitative baseline to help establish conservation priorities. PLoS One 2016 Dec 11: e0168348.

14. Sonter LJ, Ali SH, Watson JEM. Mining and biodiversity: key issues and research needs in conservation science. Proc. R. Soc. B. 2018 Dec 285: 20181926.

15. Trevelin LC, Gastauer M, Prous X, Nicácio G, Zampaulo R, Brandi I et al. Biodiversity surrogates in Amazonian iron cave ecosystems. Ecol Indicators 2019 Jun 101:813–820.

16. Trevelin LC, Simões MH, Prous X, Pietrobon T, Brandi IV, Jaffé R. Optimizing speleological monitoring efforts: insights from long-term data for tropical iron caves. PeerJ. 2021 Apr 9: e11271.

17. Umetsu F, Metzger JP, Pardini R. The importance of estimating matrix quality for modeling species distribution in complex tropical landscape: a test with Atlantic Forest small mammals. Ecography 2008 Jun 31:359–370.

18. Puttker T, Bueno AA, de Barros CD, Sommer S, Pardini R. Immigration rates in fragmented landscapes – empirical evidence for the importance of habitat amount for species persistence. PLoS ONE 2011 Nov 6:e27963.

19. Kalko EKV, Fiernel D, Handley Jr. C, Schnitzier NV. Roosting and foraging behavior of two Neotropical gleaning bats, *Tonatia silvicola* and *Trachops cirrhosus* (Phylostomidae). Biotropica 2006 Mar 34:344–353.

20. Trevelin LC, Silveira M, Port-Carvalho M, Homem DH, Cruz-Neto AP. Use of space by frugivorous bats (Chiroptera: Phyllostomidae) in a restored Atlantic Forest fragment in Brazil. Forest Ecology and Management 2013 Mar 291:136– 143.

21. Aguiar LMS, Bernard E, Machado RB. Habitat use and movements of *Glossophaga soricina* and *Lonchophylla dekeyseri* (Chiroptera: Phyllostomidae) in a Neotropical savannah. Zoologia (Curitiba) 2014 Jun 31: 223–229.

22. Kerches-Rogeri P, Niebuhr BB, Muylaert RL, Mello MAR. Individual specialization in the space use of frugivorous bats. Journal of Animal Ecol. 2020 Sep 89: 2584–2595.

23. Bernard E, Fenton B. Bat mobility and roosts in a fragmented landscape in central Amazônia, Brazil. Biotropica 2003 Mar 35:262–277.

24. Tavares, VC, Palmuti CFS, Gregorin R, Dornas TT. 2012. Morcegos. In Martins FD, Castilho AF, Campos J, Hatano FM, Rolim SG editores. Fauna da Floresta Nacional de Carajás: Estudos Sobre Vertebrados Terrestres, 161–177. Nitro Imagens, São Paulo, Brazil.

25. Handley Jr CO. 1976. Mammals of the Smithsonian Venezuelan project. Brigham Young University Science Bulletin, Provo, 20 (5): 1–91.

26. Uieda, W. 1980. Ocorrência de *Carollia castanea* na Amazônia Brasileira (Chiroptera, Phyllostomidae). Acta Amaz. 10: 936–938.

27. Tavares VC, Nobre CC, Palmuti CF, Nogueira EDP.; Gomes, J.D. et al. 2017. The bat fauna from southwestern Brazil and its affinities with the fauna of western Amazon. Acta Chiropterol. 19: 93–106.

28. Simmons, N.B., Voss, R.S., 1998. The mammals of Paracou, French Guiana, a Neotropical lowland rainforest fauna. Part 1, Bats. Bull. Am. Mus. Nat. Hist. 237: 1–219.

29. ICMBio, 2023. Sistema de Avaliação do Risco de Extinção da Biodiversidade – SALVE. Disponível em: https://salve.icmbio.gov.br/. Acesso em: 13 de nov. de 2023.

30. Gessinger, G.; Gonzalez-Terrazas, T. P.; Page, R. A.; Jung, K. & Tschapka, M. 2019. Unusual echolocation behaviour of the common sword nosed bat *Lonchorhina aurita*: An adaptation to aerial insectivory in a phyllostomid bat? Royal Society Open Science. 6: 182165.

31. Patterson, B.D. 1992. Mammals in the Royal Natural History Museum, Stockholm, collected in Brazilcand Bolivia by A.M. Olalla during 1934–1938. Fieldiana Zoology (new ser.) 66: 1–42.

32. Piló, L. B.; Auler, A. S. & Martins, F. D. 2015. Carajás National Forest: Iron ore plateaus and caves in southeastern Amazon. In: Vieira, B. C.; Salgado, A. A. R. & Santos, L. J. C.; editors. Landscapes and Landforms of Brazil. Berlin: Springer; 2015. p. 273–83.

33. Poveromo, J. J. 1999. Iron Ores The Making, Shaping, and Treating of Steel: Ironmaking Volume. Pittsburg, PA: The AISE Steel Foundation. pp. 547–550.

34. Viana, P. L. & Mota, N. F. O. 2016. Flora das cangas da Serra dos Carajás, Pará, Brasil: Styracaceae. Rodriguésia. 67: 1477–1480.

35. Devecchi, M. F.; Lovo, J.; Moro, M. F.; Andrino, C. O.; Barbosa-Silva, R. G.; Viana, P. L.; Giulietti, A. M.; Antar, G.; Watanabe, M. T. C. & Zappi, D. C. 2020. Beyond forests in the Amazon: Biogeography and floristic relationships of the Amazonian savannas. Botanical Journal of the Linnean Society. 193: 478–503.

36. MapBiomas, 2020. Projeto de Mapeamento Anual da Cobertura e Uso do Solo do Brasil. URL http://www.mapbiomas.org. Acessed in: fevereiro 2020.

37. Peel, M. C.; Finlayson, B. L. & Mcmahon, T. A. 2007. Updated world map of the Köppen-Geiger climate classification. Hydrology and earth system sciences discussions. 4: 439–473.

38. 38. Kunz, T. H.; Hodgkison, R. & Weise, C. D. 2009. Methods of capturing and handling bats. In: Kunz, TH.; Parsons, S, eds. Ecological and behavioral methods for the study of bats. Baltimore: Johns Hopkins University Press 3-35.

39. Brunet-Rossini, A. K. & Wilkinson, G. S 2009. Methods for estimating age in bats. In: Kunz, TH; Parsons, S, eds. Ecological and behavioral methods for the study of bats. Baltimore: Johns Hopkins University Press 315-326.

40. Sikes, R. S. & Gannon, W. L. 2011. Guidelines of the American Society of Mammalogists for the use of wild mammals in research. J Mammal. 92: 235–253.

41. Börger, L.; Franconi, N.; de Michele, G.; Gantz, A.; Meschi, F.; Manica, A.; Lovari, S. & Coulson, T. 2006. Effects of sampling regime on the mean and variance of home range size estimates. J. Anim. Ecol. 75: 1393–1405.

42. Jacob, A. A. & Rudran, R. 2003. Radiotelemetria em estudos populacionais. In: Cullen Jr., L.; Rudran, R. & Valladares-Padua, C. (Eds.), Métodos de Estudos em Biologia da Conservação e Manejo da Vida Silvestre. Editora da UFPR & Fundação O Boticário de Proteção à Natureza, Curitiba, pp. 285–341.

43. Rodgers, A. R.; Carr, A. P.; Beyer, H. L.; Smith, L. & Kie, J. G. 2007. HRT: Home Range Tools for ArcGIS, 1.1. Centre for Northern Forest Ecosystem Research, Ontario Ministry of Natural Resources, Ontario.

44. Kernohan, B. J.; Gitzen, R. A. & Millspaugh, J. J. 2001. Analysis of animal space use and movement. In: Millspaugh, J. J. & Marzluff, J. M. (Eds.), Radio Tracking and Animal Populations. Academic Press, San Diego, pp. 126–166.

45. White, G. C. & Garrott, R. A. 1990. Analysis of Wildlife Radiotracking Data. Academic Press, San Diego.

46. Seaman, D. E. & Powell, R. A. 1996. An evaluation of the accuracy of kernel density estimators for home range analysis. Ecology. 77: 2075–2085.

47. Johnson, D. H. 1980. The comparison of usage and availability measurements for evaluating resource preference. Ecology. 61: 65–71.

48. Fattorini, L.; Pisani, C.; Riga, F. & Zaccaroni, M. 2014. A permutation-based combination of sign tests for assessing habitat selection. Env Ecol Stat. 21:161– 187.

49. R Core Team. R Project for Statistical Computing, 2019.

50. Fattorini, L.; Pisani, C.; Riga, F.; Zaccaroni, M. 2017. The R package “phuassess” for assessing habitat selection using permutation-based combination of sign tests. Mammal Biol. 83: 64–70.

51. Zuur, A. F.; Leno, E. N. & Smith, G. M. 2007. Analysing Ecological Data. Springer, New York.

52. Burnham, K. P. & Anderson, D. R. 2002. Model Selection and Multimodel Inference: a Practical Information-theoretical Approach, 2nd edn. New York: Springer.

53. Harrison, X. A.; Donaldson, L.; Correa-Cano, M. E.; Evans, J.; Fisher, D. N.; Goodwin, C. E. D.; Robinson, B. S.; Hodgson, D. J. & Inger, R. 2018. A brief introduction to mixed effects modelling and multi-model inference in ecology. PeerJ. 6: e4794.

54. Kuznetsova, A.; Brockhoff, P. B.; Christensen, R. H. B. & Jensen, S. P. 2020. “lmerTest”: tests in linear mixed effects models, R package version 3.1–2.

55. Barton, K. 2020. MuMIn: Multi-model inference. https://CRAN.R-project.org/package=MuMIn.

56. Meyer, C. F. J.; Weinbeer, M. & Kalko, E. K. V. 2005. Home-range size and spacing patterns of *Macrophyllum macrophyllum* (Phyllostomidae) foraging over water. J. Mammal. 86: 587–598.

57. Kniowski, A. B., & Gehrt, S. D. 2014. Home range and habitat selection of the Indiana bat in an agricultural landscape. The Journal of Wildlife Management, 78(3), 503–512.

58. Lindstedt SL, Miller BJ, Buskirk SW (1986) Home range, time, and body size in mammals. Ecology 67:413–418.

59. Kelt, D.A.; Van Vuren, D. 1999. Energetic constraints and the relationship between body size and home range area in mammals. Ecology 80:337–340

60. Fenton M. B. 1997. Science and the conservation of bats. Journal of Mammalogy 78:1–14.

61. Fenton, M. B.; Whitaker Jr, J. O.; Vonhof, M. J.; Waterman, J. M., Pedro, W. A., Aguiar, L., Rautenbach, N. I. 1999. The diet of bats from Southeastern Brazil: the relation to echolocation and foraging behaviour. Revista Brasileira de Zoologia 16(4):1081–1085.

62. Falcão, F.; Ugarte-Núñez, J. A.; Faria, D.; Christini B. Caselli. (2015): Unravelling the calls of discrete hunters: acoustic structure of echolocation calls of furipterid bats (Chiroptera, Furipteridae), Bioacoustics: The International Journal of Animal Sound and its Recording 10.1080/09524622.2015.1017840.

63. Schmieder, D. A; Kingston, T.; Hashim, R; Siemers, B. M. 2010. Breaking the trade-off: rainforest bats maximize bandwidth and repetition rate of as they approach prey. Biol Lett 6(5):604–609. doi:10.1098/rsbl.2010.0114

64. Denzinger, A.; Tschapka, M. & Schnitzler H. 2018. The role of echolocation strategies for niche differentiation in bats. Canadian Journal of Zoology 96: 171– 181.

65. Ter Hofstede H. M., Ratcliffe J. M. 2016. Evolutionary escalation: the bat-moth arms race. J Exp Biol. 219:1589–602. doi: 10.1242/jeb.086686. PMID: 27252453.

66. Moir, H. M.; Jackson, J. C.; Windmill, J. F. C. 2013. Extremely high frequency sensitivity in a ‘simple’ ear. Biol. Lett.92013024120130241

67. Lassieur, S.; & Wilson, D. E. 1989. Lonchorhina aurita. Mammalian Species. 347: 1–4.

68. Goerlitz, H. R.; Greif, S. & Siemers, B. M. 2008. Cues for acoustic detection of prey: insect rustling sounds and the influence of walking substrate. J. Exp. Biol. 211: 2799–2806.

69. Weinbeer M.; Kalko E. K.; Jung, K. 2013. Behavioral flexibility of the trawling long-legged bat, *Macrophyllum macrophyllum* (Phyllostomidae). Front Physiol. 4:342. doi: 10.3389/fphys.2013.00342. PMID: 24324442; PMCID: PMC3838978.

70. Kunz T. H.; Lumsden, L. F. 2003. Ecology of cavity and foliage roosting bats. In: Kunz TH, Fenton MB (Eds) Bat Ecology. The University of Chicago Press, Chicago, 3–87.

